# Perceptions on the genetic biocontrol of invasive carp

**DOI:** 10.1101/2023.07.27.550879

**Authors:** Sam E Erickson, Pema Lhewa, Kasey Rundquist, Sayard Schultz, Lucy Levers, Michael J Smanski

## Abstract

Common carp (*Cyprinus carpio*) is an invasive species that damages native aquatic ecosystems. Advances in genetic engineering have led to novel genetic biocontrol methods that have the potential to aid in the control of carp and other invasive species. Research and development of these technologies is proceeding in parallel with outreach activities which aim to inform and engage the public. Public perceptions on the ethics, economics, and policy decisions surrounding the possible application of genetic control technologies are important in decision-making processes. Here we report results from a survey of more than 1,300 adults that measures the perceptions of risks, benefits, and overall comfort with a panel of existing and proposed carp control methods. Our population sample largely agrees that carp control is an important issue that demands action. We observe trends suggesting that comfort levels with the possible future implementation of genetic biocontrol for common carp are strongly influenced by the perception of benefits compared to other methods, in particular effectiveness and environmental safety. No statistically significant correlations between comfort levels and river/lake use habits nor demographics are observed. There is a need for further research to identify how respondents from similar backgrounds come to different knowledge and attitudes on the matter.

## Introduction

Common carp (*Cyprinus carpio*) is an invasive pest that poses a significant threat to US water quality and freshwater ecosystems. Carp thrive in shallow lakes, rivers, and marshes, and since being introduced in the United States in the late 1800’s they have spread throughout the country (1). Many lakes and rivers in North America have reached carp densities up to 1,000 kg body mass per hectare (2). Bottom feeding of invasive carp results in increased water turbidity, release of excess nutrients from sediments, and destruction of vegetation. Turbid water prevents sunlight from passing through lake surfaces and also inhibits the growth of native game fish that hunt by sight. Vegetation loss threatens waterfowl that are unable to thrive in carp-dominated lakes. Lakes and rivers with large carp populations lose their ability to maintain water clarity by trapping and transforming nutrients, which accelerates the problem of waterway eutrophication from agricultural run-off (3). Many of these negative impacts can be prevented by maintaining carp densities below a management target level of 100 kg/ha (4). Carp are exceptionally difficult to control, as carp reproduce rapidly, with females laying up to 2 million eggs in a single spawning period. As such, there is a demand for new and improved methods for population control of common carp.

Novel genetic biocontrol technologies may pose a viable solution for carp control in the near future. Genetic biocontrol refers to a range of new pest control methods made possible by genetic engineering, in which a pest organism is converted into a pesticide agent. Genetic biocontrol can take many forms, such as the wild release of organisms that alter the sex-ratio of future generations or that disrupt normal reproduction needed to sustain population size (5). Genetic biocontrol of common carp first came to be recognized as a feasible option in the late 1990s (6). The Daughterless Carp Project, which sought to genetically modify carp such that they only produce male offspring, served to spread awareness and initiate discussion through public engagement (7). Since then, advances in genetic engineering have made possible new genetic bio-control strategies. Tools such as CRISPR/Cas9, which enables precision gene modification, serve to accelerate the progression from design to implementation of genetically engineered systems. The progress has generated cautious optimism regarding its potential uses in conservation biology (8). In parallel to this technology development, attitudes surrounding genetic biocontrol have evolved. In a 1987 publication on public perceptions of risk, DNA technology was seen as one of the most risky of 81 diverse technologies (9). It was an outlier in two metrics of risk: the potential risks were seen both as dreadful and unknown. A 2004 survey on approaches for control of aquatic invasive species reported that, among 12 management options deemed acceptable under some circumstances (3 options were deemed unacceptable under any circumstance), genetic manipulation of the invasive species ranked lowest in acceptability (10). In 2011, focus group studies of stakeholders in aquatic systems management identified numerous benefits of genetic biocontrol, but still noted that we were still far from public acceptance (11). These focus groups cited ecological risks, unknown risks, and financial risks as top concerns.

The past decade has witnessed an increase in public acceptance of genetic biocontrol methods as a viable option. For example, a 2018 survey of 2500 Americans by the Pew Research Institute found a 7:3 ratio of support over opposition for engineering the genome of mosquitoes to limit their reproduction and slow the spread of disease (12). In a 2019 survey of the Great Lakes region of the north central USA, 85% of professional stakeholders and 95% of the fishing community supported or strongly supported the research and development of genetic biocontrol to combat sea lampreys in the Great Lakes (13). The same year, a survey of 1600 US adults led by the University of Wisconsin found that the perceived benefits outweighed perceived risks for applications of genome editing in conservation biology, even though specific examples of how genome editing would be applied were not included (14). In a survey covering the use of gene drives in agriculture, most of the >1000 respondents supported suppression or replacement gene drives as long as there were built-in controls to prevent their spread (15). Despite this overall increase in support for genetic biocontrol, local opinions can vary. In a classic example of ‘not in my backyard’ (NIMBY), a 2016 local vote on a proposed GE mosquito field trial in southern Florida, the county residents voted in support while the residents of the city in which the trial would take place voted in opposition.

Although genetic biocontrol technologies for carp are still in development, we believe that now is the time for public outreach efforts in order to better understand public opinion on the matter. We present here a local survey of Minnesota residents asking for their perceptions regarding options for invasive carp population control, genetic biocontrol, as well as collecting information on demographics and river/lake use habits. Schairer et al. recently published a typology for public engagement efforts concerning genetic biocontrol (16). In this model, public engagement falls into one of three categories; engagement to inquire, engagement to involve, or engagement to influence. Our study falls under the category of engagement to inquire. We did not seek to involve the public in decision making at this point, and we took efforts to avoid influencing survey respondents, as noted below. We hypothesized that the comfort level of Minnesota residents with respect to the possible future use of genetic biocontrol for carp management may correlate with demographics or river/lake use habits.

Our population sample largely recognizes controlling carp populations as an important issue which deserves intensive measures, and expressed awareness of a variety of methods. Our correlation analysis revealed no strong correlation between comfort level with genetic biocontrol and demographic data nor river/lake use habits. Comfort with the use of genetic biocontrol seems to be heavily influenced by perceptions of particular benefits to such approaches, in particular efficacy when compared to existing methods, and lower risk of harm to native fish. We see a need for further research to assess why people from similar demographics and with similar water use habits come to different views on genetic biocontrol.

## Methods

### Survey Design

The survey was organized in four general sections: (i) use of Minnesota lakes/rivers; (ii) attitude towards physical, chemical, and biological management options for invasive carp; (iii) attitude towards genetic biocontrol methods for invasive carp; and (iv) demographics.

Section (i) includes general questions relating to how often respondents use Minnesota’s lakes, how they use lakes, observed changes in the conditions of the lakes, and attitudes regarding environmental and water quality.

Section (ii) begins with background information that provides some context for common carp as an invasive species. The section then includes questions aimed towards the importance of controlling carp species in Minnesotan lakes, how strictly they should be controlled, and who should be responsible for the control. Next, there are a series of questions asking respondents about their familiarity with myriad control options, the perceived effectiveness of each control option, the likelihood of each control option to harm native fish populations, and their overall comfort level with each option. Section (iii) begins with a definition of specific genetic biocontrol methods including sterile-male technology, sex-ratio-biasing technology, and gene drive technology. Following these definitions, respondents were then asked about their likelihood to support the use of each technology, the possible risks and benefits (wherein many potential risks and benefits were presented as a list from which the respondents could select any that applied), and what information they would want to know before deciding whether or not to support a technology.

Section (iv) collects demographic information about the survey respondent, including gender, age, ethnicity, education, and income.

Following feedback from presenting results of the initial 2018 email survey to local stakeholders, we made a slight modification to the survey prior to administration at the Minnesota State Fair. We replaced the word ‘poison’ from the description of chemical control with ‘chemical that kills fish’. Both versions (*i*.*e*. with and without this language replacement) were administered at the Minnesota State Fair in equal numbers to control for the possibility that the two survey populations (email versus state fair) would respond differently to these questions irrespective of the language used.

### Data collection

The survey was administered to adult (18 and older) participants during the Fall of 2018 or Summer of 2019. Prior to administration, this study was determined exempt by the University of Minnesota (UMN) Institutional Review Board. The survey was administered in two ways. It was first emailed to a list-serve for the Minnesota Aquatic Invasive Species Research Center (approximately 3,100 recipients), from which 603 responses were collected. This list-serve includes stakeholders in watershed management, natural resources preservation, and many ‘Aquatic Invasive Species Detectors’ who report observations of aquatic invasive species. The second administration was in-person at the 2019 Minnesota State Fair (Driven to Discover Building) where 703 people took the survey on a touchscreen tablet. Respondents were able to skip any question in the survey, but this was rear in our experience. For the in-person State Fair surveying, staff were instructed not to discuss research of genetic biocontrol with the survey respondents to avoid biasing the responses(17). Participation in the State Fair Survey was incentivized by free drawstring tote-bags. A total of 1306 responses were collected using Google forms. Survey results for all respondents, a codebook explaining how data is encoded, a summary of all variable counts, as well as a copy of the text of the complete survey are included in the supplement. In our data set, categorical ordinal variables are integer-encoded, and binary variables are dummy-encoded as described in the accompanying codebook.

### Data Analysis and Statistics

We conducted our analysis in three parts. First, we report a summary of the demographic distribution of our population sample, as well as their self-reported lake/river water use habits. This information helps characterize our population sample, and assist in making judgements regarding the applicability of results to other communities. Secondly, we report a summary of survey responses regarding respondent knowledge and opinion towards control of invasive carp. Finally, we assess correlations between respondent level of comfort with the possible future implementation of genetic biocontrol in Minnesota and other survey responses, such as demographics, lake and river use habits, and attitudes towards invasive species management and genetic biocontrol methods.

Our key outcome variable was perceived level of comfort with the possible future implementation of genetic biocontrol in Minnesota. Level of comfort was defined on a scale, with possible responses of “very comfortable”, “moderately comfortable”, “uncomfortable”, “very uncomfortable”, or “I don’t know”. These options were presented from left to right across the screen, and only a single box could be checked (e.g., for each control option only one of the comfort levels could be selected). For correlation analysis, we calculated Spearman’s ρ (also known as Spearman’s rank correlation coefficient) for our key outcome variable and 102 survey response predictor variables. Spearman’s ρ is commonly used for analysis correlation between survey data of this format, as it reports a nonparametric measure of rank correlation which doesn’t pre-suppose any particular statistical distribution, and therefore is appropriate for non-linear relationships, for instance between ordinal categorical variables. For correlation analysis, we apply a Bonferroni-corrected threshold for significance of p<0.0005. Since we conducted 102 statistical tests in parallel, we divide a conventional p value threshold of 0.05 by 102. For ordinal categorical variables, missing data and noncommittal answers such as “I don’t know” were not included in correlation analysis.

## Results

### Demographic composition of survey respondents

When both surveying methods are combined, we received 1306 responses. The demographics of survey takers are reported in Table 1. Residents from across the state of Minnesota participated, with the greatest concentration of participants residing in zipcodes near the Minnesota State Fair, which took place the city of Saint Paul in zipcode 55108 (Fig. S1, S2). Our population sample included an over-representation of male-identifying respondents, with 704 male and 535 female. The numbers of respondents who self-identified as white (83%) versus non-white approximates the general population of Minnesota (85% white), although the specific distribution of minority races/ethnicities does not precisely represent those distributions among the Minnesota public (18). The highest educational degree attained by survey respondents was substantially higher than expected for the overall population. The most recent US Census data report 36% of the national population have at least a Bachelor’s degree, while 82% of our respondents had a Bachelor’s degree or higher (18). Our population sample includes many habitual users of Minnesota lakes and rivers. A majority of respondents reported frequent (>4 times per month, 43%) or moderate use (1-4 times per month, 23%), with 29% reporting infrequent (<1 time per month) and the rest reporting that they have never used Minnesota lakes or rivers. The most common uses were for swimming (n=800), fishing (n=799), and non-motorized boating (n=658). A majority of respondents thought that Minnesota lakes and rivers are in good condition (n=784). Interestingly, 66% of respondents reported that they have noticed a change in the conditions of Minnesota lakes and rivers, and among those respondents, 83% reported noticing a worsening of the condition of Minnesota lakes and rivers. Respondents who reported specific types of lake/river use (*i*.*e*. fishing, hunting (non-)motor boat usage, and swimming) were more likely to observe that the conditions of Minnesota water resources are worsening (Pearson’s Chi-squared test: x^2^(1,N=1299) = 19, p < 10^−4^). Among respondents who reported specific types of lake/river use, 57% observed that conditions of Minnesota water resources are worsening, as opposed to 37% among respondents who did not report specific lake/river use habits.

**Table 1.** Demographics of survey respondents. Each topic had several respondents who elected not to answer.

| Topic | Selection | Email Survey | State Fair Survey | Total |
| --- | --- | --- | --- | --- |
| Sex | Female | 174 (28%) | 361 (51%) | 535 (40%) |
|  | Male | 402 (66%) | 302 (42%) | 704 (53%) |
| Race | Asian | 9 (1%) | 30 (4%) | 39 (2%) |
|  | Black or African American | 5 (<1%) | 22 (3%) | 27 (2%) |
|  | Native Hawaiian/Pacific Islander | 2 (<1%) | 3 (<1%) | 5 (<1%) |
|  | Hispanic | 6 (<1%) | 22 (3%) | 28 (2%) |
|  | Indigenous American | 8 (1%) | 13 (2%) | 21 (1%) |
|  | White | 526 (87%) | 587 (83%) | 1113 (85%) |
|  | No answer | 60 (10%) | 59 (8%) | 119 (9%) |
| Age | 18-20 years | 2 (<1%) | 39 (5%) | 41 (3%) |
|  | 21-30 years | 66 (10%) | 98 (13%) | 164 (12%) |
|  | 31-40 years | 96 (15%) | 87 (12%) | 183 (14%) |
|  | 41-50 years | 64 (10%) | 91 (12%) | 155 (11%) |
|  | 51-60 years | 101 (16%) | 151 (21%) | 252 (19%) |
|  | 61-70 years | 159 (26%) | 152 (21%) | 311 (23%) |
|  | >70 years | 83 (13%) | 56 (7%) | 139 (10%) |
| Highest Degree | Less than a high school diploma | 1 (<1%) | 8 (1%) | 9 (<1%) |
|  | High School, or equivalent | 33 (5%) | 132 (18%) | 165 (12%) |
|  | Bachelor's | 266 (44%) | 346 (49%) | 612 (46%) |
|  | Master's | 198 (32%) | 123 (17%) | 321 (24%) |
|  | Doctoral | 80 (13%) | 60 (8%) | 140 (10%) |
| Yearly household income | <\$20,000 | 10 (1%) | 40 (5%) | 50 (3%) |
| | \$20,000-\$34,999 | 19 (3%) | 51 (7%) | 70 (4%) |
| | \$35,000-\$49,999 | 42 (6%) | 63 (8%) | 105 (8%) |
| | \$50,000-\$74,999 | 102 (17%) | 98 (13%) | 200 (15%) |
| | \$75,000-\$99,999 | 113 (18%) | 88 (12%) | 201 (15%) |
| | >\$100,000 | 203 (33%) | 224 (31%) | 427 (32%) |
| Total |  | 603 | 703 | 1306 |

Overall, the 1306 survey respondents in this study are not statistically representative of the general population. They are slightly enriched for males and substantially enriched for college and graduate degree attainment. Despite this, the responses surrounding lake and river use frequency and mode show that we have collected the perspectives of a diverse set of stakeholders from across Minnesota.

### Awareness of common carp population control strategies

Our sample population largely views carp control as an important issue. Following a brief paragraph describing the impact of invasive carp on lake ecosystems (see survey text in Supplementary Information), survey respondents were asked several questions regarding population control strategies. Respondents were asked to rate the importance of decreasing the impact of carp on Minnesota lakes and rivers on an ordinal scale (very important, important, neutral, not important, unknown). Over 80% of respondents considered decreasing the impact of carp on Minnesota lakes/rivers important or very important (Fig. 1a).

**Fig. 1.**
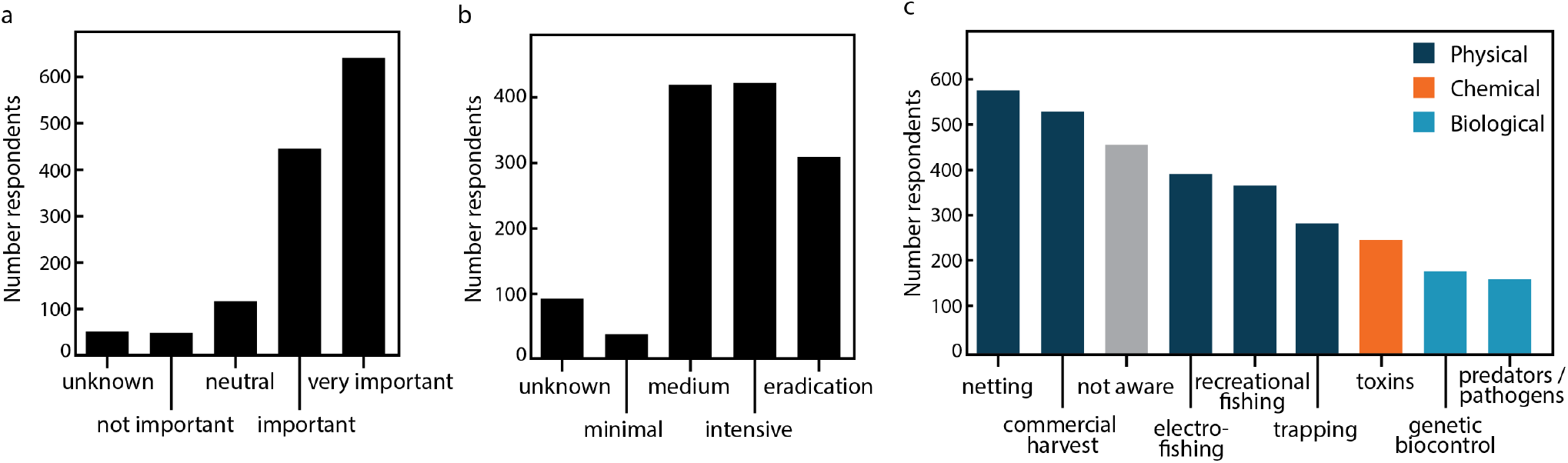
General awareness of carp control. (**a**) Survey responses regarding the level of importance of controlling invasive common carp. (**b**) Opinions towards the rigor of carp control that should be implemented. Multi-choice response options included definitions: eradication control (removal of all carp in a lake), intensive control (removal of most carp in a lake), medium control (removal of just enough carp to prevent ecosystem damage), minimal control (decrease the carp numbers, but not enough to limit ecosystem damage), and unknown. (**c**) Awareness of different methods for carp control, color coded by physical methods (dark blue), chemical methods (orange), biological methods (light blue), or respondents that were not aware of any control methods (grey). Survey respondents could select multiple answers for the question captured by panel (**c**).

Respondents did not largely agree as to the level of intensity of population control that should be implemented. The subset of respondents who selected “yes” or “maybe” regarding whether action should be taken to control carp populations were asked how strictly common carp should be controlled. Answers to the multiple-choice question followed an ordinal scale: “Eradication control” (removal of all carp in a lake), “Intensive Control” (removal of most carp in a lake), “Medium control” (removal of just enough carp to prevent ecosystem damage), “Minimal control” (decrease the carp numbers, but not enough to limit ecosystem damage), and “Unknown”. Responses were split fairly evenly between medium levels of control, intense control, and eradication (Fig. 1b). Respondents were allowed to choose multiple responses regarding who should be responsible for carp population control, and the most frequent response was the government (n = 1072), followed by lake associations (n = 742), private companies (n = 439), and then individual citizens (n = 436).

Respondents were presented a list of carp control methods, and asked to identify which ones they were aware as existing. For Minnesota lakes and rivers managed by the Minnesota Department of Natural Resources, commercial seining (used in 19 lakes) and box netting (17 lakes) are the most commonly used carp removal techniques, followed by drawing down water levels (11 lakes) or using rotenone for chemical control (9 lakes). This roughly parallels the familiarity with different techniques reported by the survey respondents, who were more likely to be aware of commercial harvest and netting and less likely to be aware of chemical and biocontrol options (Fig. 1c). In general, awareness of physical methods for carp control were more recognized than chemical or biological control options. Most respondents (82%) believe that the government should be responsible for carp control efforts. This mirrors a recent survey conducted on carp biocontrol in Australia, where there was a further preference for placing decision-making power in local governments over state and federal agencies (19).

### Perceptions of physical, chemical, and biological control of aquatic invasive species

When asked about their comfort level with the various population control approaches, a clear pattern emerged. Survey takers were asked to express their level of comfort with the implementation of various control methods on an ordinal scale (Very comfortable, Moderately comfortable, Uncomfortable, Very Uncomfortable, and I don’t know). When approaches were grouped by category, respondents showed the most comfort with physical methods, followed by biological methods, and finally chemical methods (Figure 2a). Of the biological methods, genetic biocontrol received the highest comfort scores but also the largest number of “I don’t know” responses. Release of pathogens received the lowest comfort scores of the biological approaches. Chemical methods received the lowest scores for comfort level regardless of whether the respondents received a survey version describing chemical agents as “poisons” or as “chemicals that kill fish”.

**Fig. 2.**
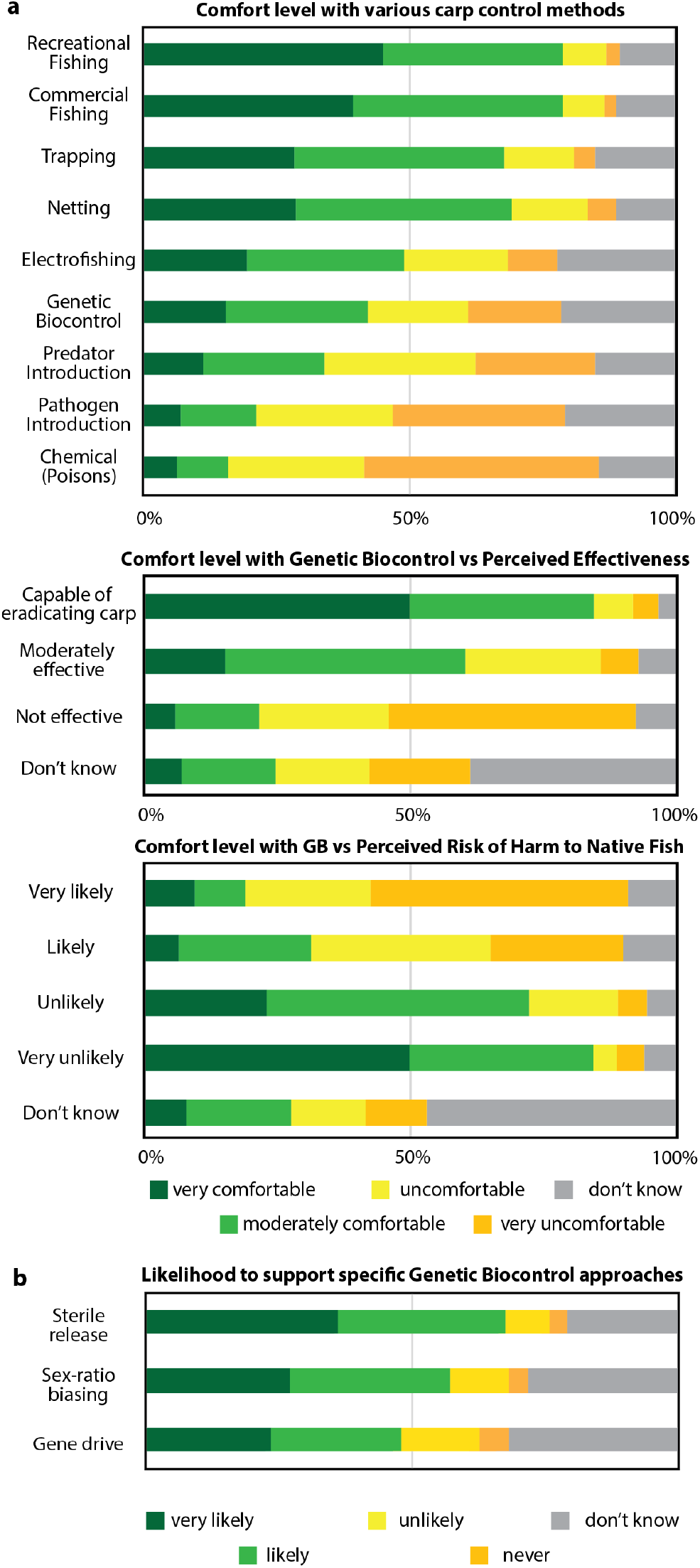
Comfort with or support of different carp control strategies. (**a**) Horizontal bar graph showing the fraction of reported comfort levels with general carp control approaches (top), or specifically towards Genetic Biocontrol (middle and bottom) for specific subgroups of respondents. In the middle plot, responses were divided based on how respondents answered a different question pertaining to the perceived effectiveness of genetic biocontrol (GB) compared to traditional approaches. In the bottom plot, responses were divided between respondents based on how respondents answered a different question pertaining to the perceived likelihood of harm to native fish resulting from genetic biocontrol. (**b**) Self-reported likelihood to support specific genetic biocontrol approaches.

The third part of the survey sought to gather opinions regarding specific genetic biocontrol strategies. The survey text provided a short, 2-3 sentence description of three genetic biocontrol strategies: sterile release, sex-ratio biasing, and suppression gene drive. Respondents were instructed to read those descriptions before reporting their likelihood to support the use of the methods using an ordinal scale (Very likely, Likely, Unlikely, Never, and I need more information). Of the three methods, respondents expressed the greatest likelihood to support sterile release, followed closely by sex ratio biasing (Figure 2b). Respondents expressed the least likelihood to support the use of gene drives, which was also the method which received the greatest number of “I don’t know” responses.

### Notable correlations in survey results

For this study, self-reported comfort levels with the implementation of genetic biocontrol methods in Minnesota represents a key outcome variable. The ability to predict comfort levels based on demographic variables, water use habits, or opinions on other subjects would be useful to policy makers and wildlife managers, as it would help reveal the forces that drive public opinion and help with allocating resources in public outreach efforts. We performed a correlation analysis to determine which survey responses have predictive power with respect to comfort levels for genetic biocontrol. For all of the binary-encoded and ordinal categorical-encoded survey questions, we calculated Spearman’s rank correlation coefficient (Spearman’s ρ) and p-values, as shown in Supplementary Table S1. Since we are simultaneously testing 102 possible bivariate relationships, we apply a Bonferroni-corrected p value threshold of p<0.0005 (0.05/102) for significance.

In general, we find that demographics and lake/river use habits poorly predict comfort with the use of genetic biocontrol. We observed no strong correlations between comfort levels with genetic biocontrol methods and responses to any question in survey section 1/4: Use of Minnesota Lakes and Rivers nor section 4/4: Demographics (Table S1, S4). As such, although perhaps subgroups of our population sample may have shared similar attitudes towards carp control and possible technologies, it appears we may not be able to accurately predict where people are in the distribution based on demographics and water-use habit information alone.

Comfort with the use of genetic biocontrol appears to be influenced by whether respondents see it as having specific benefits. Our population sample seems roughly evenly divided as to whether genetic biocontrol poses any meaningful advantages. Participants were asked “What are the benefits of Genetic bio-control strategies?”, and prompted to select one or more answers from a list of possible benefits. An answer of “I don’t know” strongly anticorrelated with genetic bio-control comfort level l (Spearman’s ρ = −0.3, p = 1.4×10^−21^). Approximately half, 54%, (n=673) of our sample identified one or more benefit to genetic biocontrol. The list of possible benefits included “They are more effective than available control methods”, “They are less costly than available control methods”, “They are safer to humans than available control methods”, “They are safer for the environment than available control methods”, and “They have less risks compared to other control methods”. Among this subset, 61% self-reported as “moderately comfortable” or “very comfortable” with genetic biocontrol. Among those who did not identify any benefits, this proportion drops to 20%. The most commonly identified benefits were environmental safety (36% of all respondents) and comparative effectiveness (33% of all respondents).

It appears that perceptions of efficacy may be a strong driver of comfort with the use of genetic biocontrol (Figure 2a). Survey respondents were asked were asked to rank their perceived effectiveness of several carp population control methods on an ordinal categorical scale, with possible answers “Capable of eradicating invasive carp”, “Moderately effective”, “Not effective”, and “I don’t know.” “I don’t know,” and missing answers were not included in correlation analysis. Survey respondents who perceived genetic biocontrol as more effective than other methods were much more likely to report higher levels of comfort with its implementation in Minnesota (Spearman’s ρ = 0.53, p = 3.3×10^−45^, table S2).

Perception of risk for harm to native fish populations appears to be a strong driver of discomfort with genetic biocontrol (Figure 2a). Survey respondents were asked were asked to estimate the likelihood of harm to harm to native fish populations associated with several carp population control methods on an ordinal categorical scale, with possible answers “very likely”, “likely”, “unlikely”, “very unlikely” (“I don’t know” or declining to answer were options, but excluded from correlation analysis). Perception of a high likelihood of harm from genetic biocontrol inversely correlated with comfort levels (Spearman’s ρ = −0.55, p = 1.7×10^−57^, table S2).

## Discussion

This study aimed to evaluate the public perceptions in Minnesota surrounding possible future genetic biocontrol of invasive carp. Although this technology is not yet mature and still in development, now is the time to start seriously considering the ethical and societal ramifications. We report a preliminary exploratory study seeking to form a general base of knowledge from which to formulate more specific hypotheses for future rigorous testing. In our report we include a summary of the demographics of our sample population, statistics regarding awareness and attitudes towards invasive carp and population control options, and a correlation analysis comparing comfort with genetic biocontrol of carp to other survey answers. Our hope is that studies such as this can help to identify key predictor variables that influence public knowledge and opinion towards carp biocontrol and other carp control methods. Such knowledge can help local political leaders, policy makers, and wildlife managers to more efficiently allocate time, effort, and resources towards public outreach projects.

Overall, there is a high level of awareness of the problems posed by invasive carp in our sample population, and this is seen as a very important problem (Figure 1a) that should be addressed. However, respondents did not largely agree to how intensively the population should be controlled (Figure 1b). Among status quo control methods, respondents expressed greater comfort with physical over chemical or biological methods (Figure 2a).

Our correlation analysis suggests that key drivers of attitudes towards genetic biocontrol include perceptions of specific advantages. Comfort levels correlated strongly with whether or not respondents identified genetic biocontrol has having specific benefits setting it apart from other methods. Approximately one half of our sample identified no specific benefits associated with genetic biocontrol. In particular, comfort with genetic biocontrol correlated with perceptions that it may be more effective or more environmentally safe than other methods (Figure 2a). The importance of perceived effectiveness is in line with Rogers’ theories on technology adoption, which list the ‘relative advantage’ of a new technology over competing technologies as the first of five technology attributes that drive the speed of adoption (20). Our results suggest that future public outreach related to genetic biocontrol may benefit from a focus on clearly explaining specific advantages of such technologies over existing methods.

Although we hypothesized that comfort levels with the possible future implementation of genetic biocontrol of carp in Minnesota would correlate with the demographics and lake/river use habits of our survey respondents, in our sample size of 1306 respondents, we observed no strong correlations of this type. Respondents belonging to similar demographics and with similar water use habits frequently have dramatically different attitudes towards genetic biocontrol. The lack of correlations with demographic information is congruent with that observed in a recent survey in New Zealand, where demographics, political affiliation, and religiosity had less predictive power regarding attitudes towards gene-drives than did general world-view(21). We did not ask questions that would indicate respondents’ world view, their trust in science, nor their general comfort level with new technology. These would be interesting to probe in future efforts. It appears that existing data on Minnesota communities may have limited power to predict to predict attitudes towards genetic biocontrol. Therefore, we see a need for more research into the factors which would influence people of similar demographics and with similar water use habits to come to very different attitudes.

The survey was administered through multiple platforms to mitigate biases that could be present in a single sample (22). It was first distributed via email list-serve by the Minnesota Aquatic Invasive Species Research Center (MAISRC) (approximately 3,100 recipients) from which more than 600 responses were collected. This list-serve includes stakeholders in watershed management, natural resources preservation, and certified ‘AIS Detectors’ who report observations of aquatic invasive species. The second administration was in-person at the 2019 Minnesota State Fair where more than 700 people took the survey on a touchscreen tablet. Several of the demographics questions revealed that our sample of respondents is not statistically representative of the broader population (e.g., higher educational degree attainment than the general population), and the results of this survey should be interpreted with that in mind. Our population sample exhibited greater awareness of carp as an invasive species than other surveys of invasive species or aquatic invasive species(23, 24). Eiswerth et al. reports a statistically significant correlation between completing a college degree and AIS awareness(23). Our sample was disproportionately weighted towards college degree attainment, although we observed no significant correlation between awareness of invasive carp and degree attainment.

Our survey was designed to mitigate framing bias. When an individual has no strong prior opinion regarding an issue, they are often more easily influenced by positive or negative framing (25). The paragraphs provided at the start of survey section 3 describing how each genetic biocontrol technology works can be characterized as a neutral-to-positive framing. They do not mention the risks or inherent failure modes of each technique (i.e., negative framing), but they also do not present the techniques as silver-bullet solutions to pest control (i.e., positive framing). Negative framing has been shown to produce a more lasting influence than positive framing, but when people are provide competing viewpoints, the most recent framing typically has the dominant effect (25). We expect that additional study of framing bias on attitudes towards genetic biocontrol could be informative.

Watershed and fisheries managers need to consider not only effectiveness and risk, but also public perceptions when making management decisions. With this study, we aim to better understand public perception in Minnesota towards genetic biocontrol of invasive carp. Although we observed no strong correlations between survey responses and collected demographic and lake/river use habit information, we see strong correlations between respondent comfort with the possible future implementation of genetic biocontrol with carp and perceptions of unique advantages of such methods. Insights from this study and further investigation provide a knowledge base to inform better decision making in the service of communities and ecosystems.

## Statements and Declarations

## ACKNOWLEDGEMENTS

We thank Fred Gould (Genetic Engineering and Society Center, NCSU) for helpful insights when discussing this survey data. This project was funded by a grant from the Minnesota Aquatic Invasive Species Research Center through the Environment and Natural Resource Trust Fund. The authors have no relevant financial or non-financial interests to disclose.

## AUTHOR CONTRIBUTIONS

MJS and SS designed the survey. MJS, SEE, RK, and LL collected data. MJS, SEE, and PL performed data analysis. MJS wrote the manuscript. All authors revised the manuscript.

## DATA AVAILABILITY

All data associated with this survey is available upon request. Please contact the corresponding author for data requests.

---

**Supplemental Table S1.** Spearman correlations for section 1 of survey (lake/river use) to ‘comfort with genetic biocontrol’.

| Predictor Variable | Spearman's $\rho$ | p value | Significance | Variable label (type of variable) |
| --- | --- | --- | --- | --- |
| location | -0.040 | 2.11E-01 |  | survey administered by email or MN State Fair (binary) |
| use | 0.004 | 8.94E-01 |  | frequency of use of MN lakes, rivers (ordinal categorical) |
| fishing | -0.010 | 7.48E-01 |  | used MN lakes, rivers for fishing in the past year (binary) |
| hunting | 0.042 | 1.84E-01 |  | used MN lakes, rivers for hunting in the past year (binary) |
| motor | -0.013 | 6.87E-01 |  | used MN lakes, rivers for boating (motorized) in the past year (binary) |
| nonMotor | -0.012 | 7.10E-01 |  | used MN lakes, rivers for boating (non-motorized) in the past year (binary) |
| swimming | 0.006 | 8.50E-01 |  | used MN lakes, rivers for swimming in the past year (binary) |
| nonUser | -0.047 | 1.45E-01 |  | did not use MN lakes, river in the past year (binary) |
| other | -0.005 | 8.88E-01 |  | used MN lakes, rivers for other purpose in the past year (binary) |
| goodCond | -0.010 | 7.54E-01 |  | noticed condition of MN lakes, rivers getting better (binary) |
| change | -0.010 | 7.58E-01 |  | noticed change in condition of MN lakes, rivers (ordinal categorical) |
| quality | -0.057 | 7.53E-02 |  | observed feature of MN lakes, rivers not in good condition—water quality (binary) |
| stability | -0.038 | 2.36E-01 |  | observed feature of MN lakes, rivers not in good condition—bank stability (binary) |
| clarity | -0.007 | 8.29E-01 |  | observed feature of MN lakes, rivers not in good condition—water clarity (binary) |
| Birdlife | 0.016 | 6.16E-01 |  | observed feature of MN lakes, rivers not in good condition—Birdlife (binary) |
| nativeFish | 0.012 | 7.04E-01 |  | observed feature of MN lakes, rivers not in good condition—Native fish abundance, size (binary) |
| nativeVeg | 0.006 | 8.56E-01 |  | observed feature of MN lakes, rivers not in good condition—Native vegetation abundance (binary) |
| invVeg | 0.067 | 3.51E-02 |  | observed feature of MN lakes, rivers not in good condition—Amount of non-native (invasive) vegetation (binary) |
| invFish | 0.090 | 4.89E-03 |  | observed feature of MN lakes, rivers not in good condition—Amount of non-native (invasive) fish (binary) |
| Algal | 0.012 | 6.99E-01 |  | observed feature of MN lakes, rivers not in good condition—Algal blooms (binary) |
| NoneEvery | -0.013 | 6.80E-01 |  | observed that every feature of MN lakes, rivers are in good condition (binary) |

**Supplemental Table S2.**
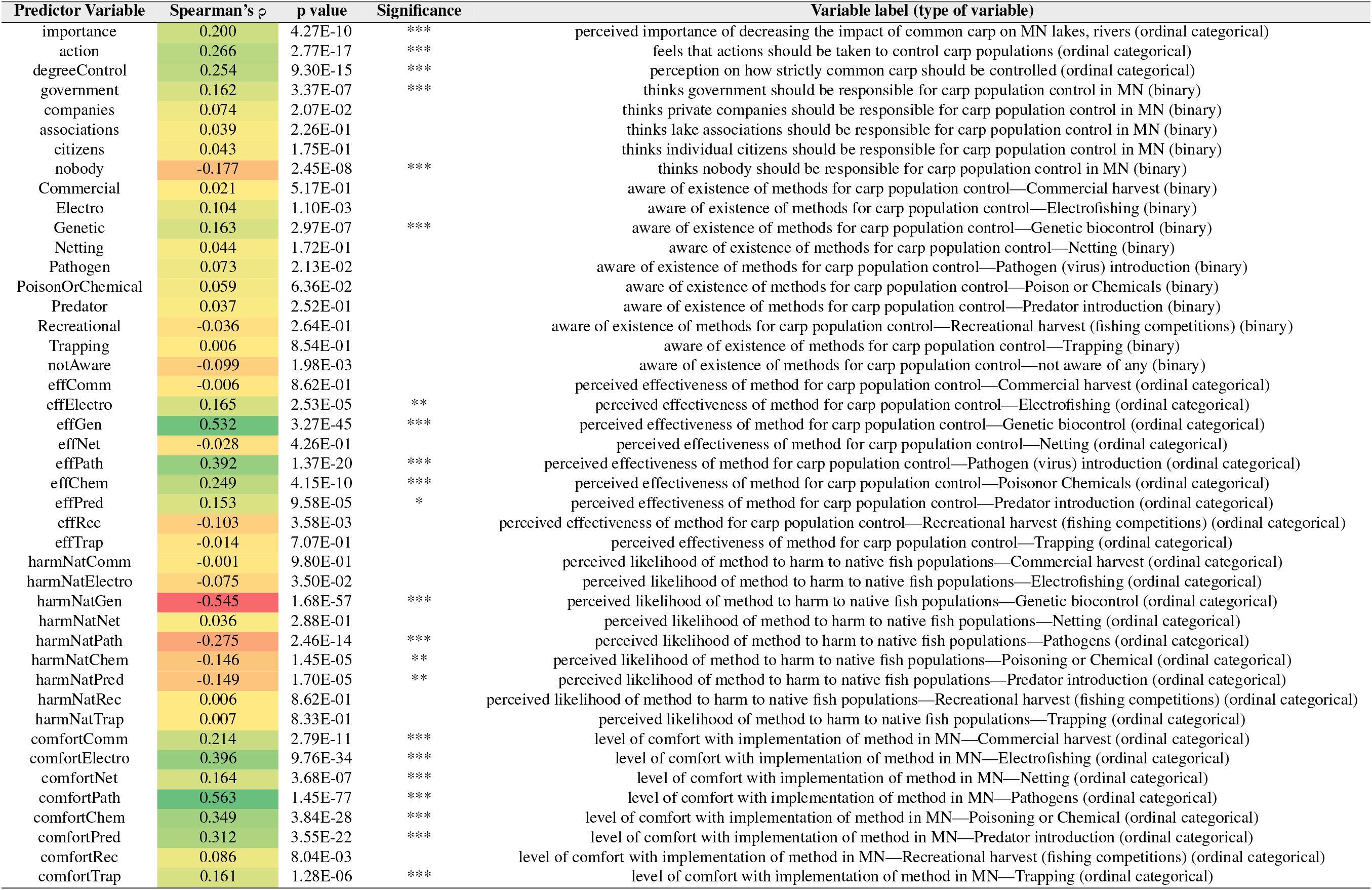
Spearman correlations for section 2 of survey (attitude towards management options) to ‘comfort with genetic biocontrol’.

**Supplemental Table S3.**
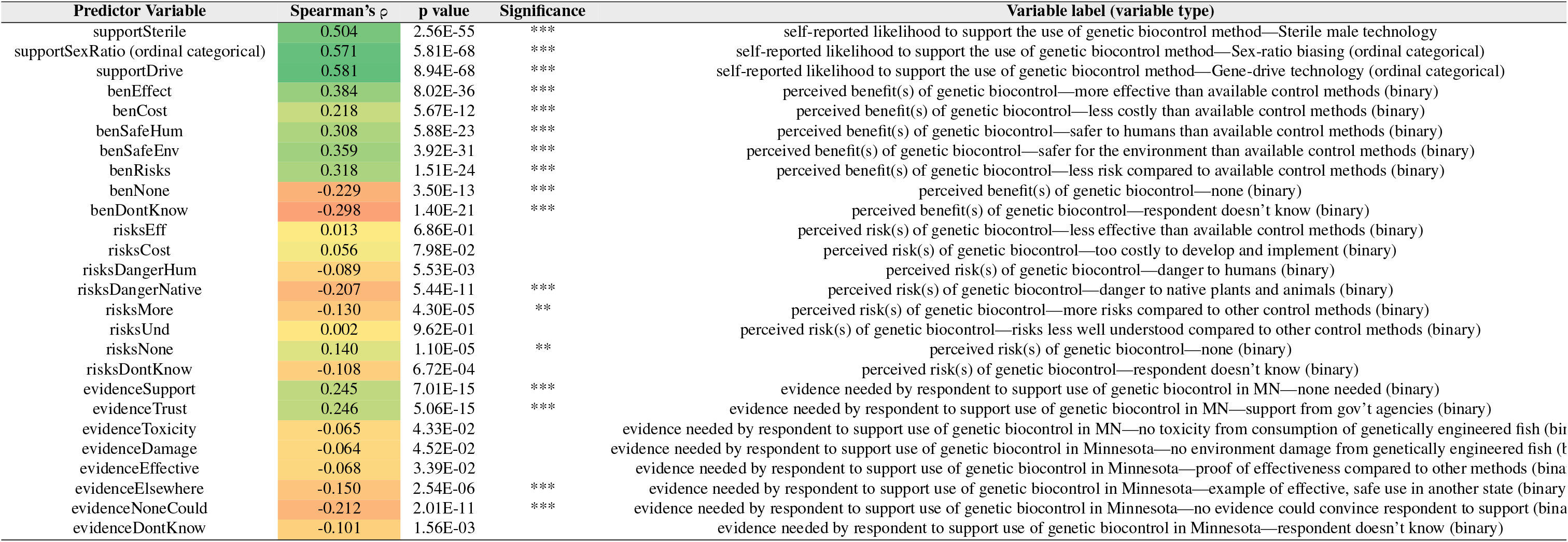
Spearman correlations for section 3 of survey (genetic biocontrol methods) to ‘comfort with genetic biocontrol’.

**Supplemental Table S4.**
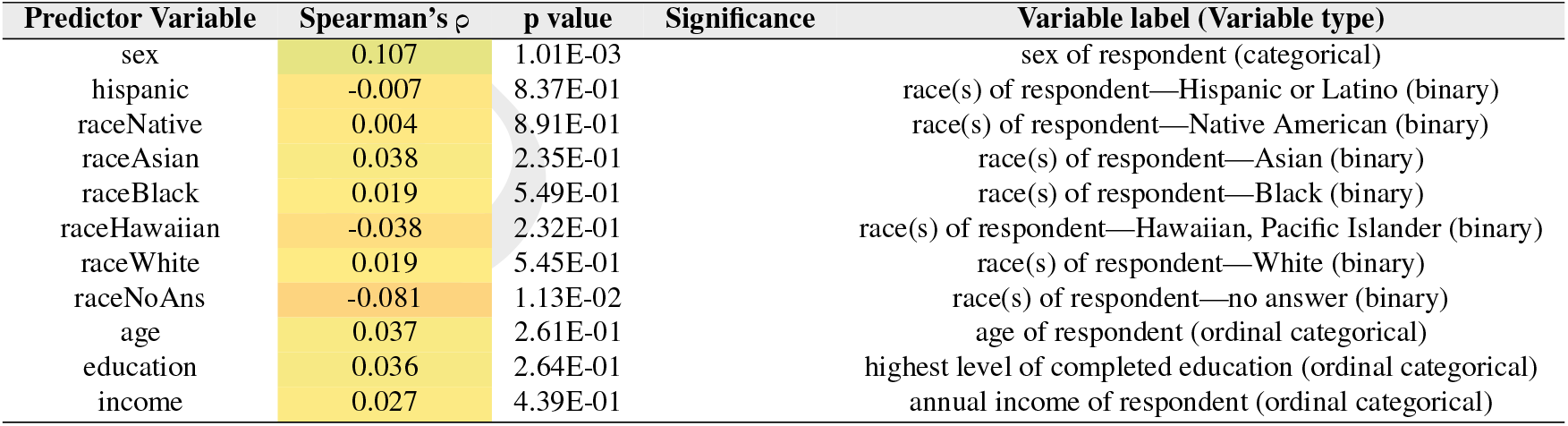
Spearman correlations for section 4 of survey (demographic information) to ‘comfort with genetic biocontrol’.

**Supplemental Figure S1.**
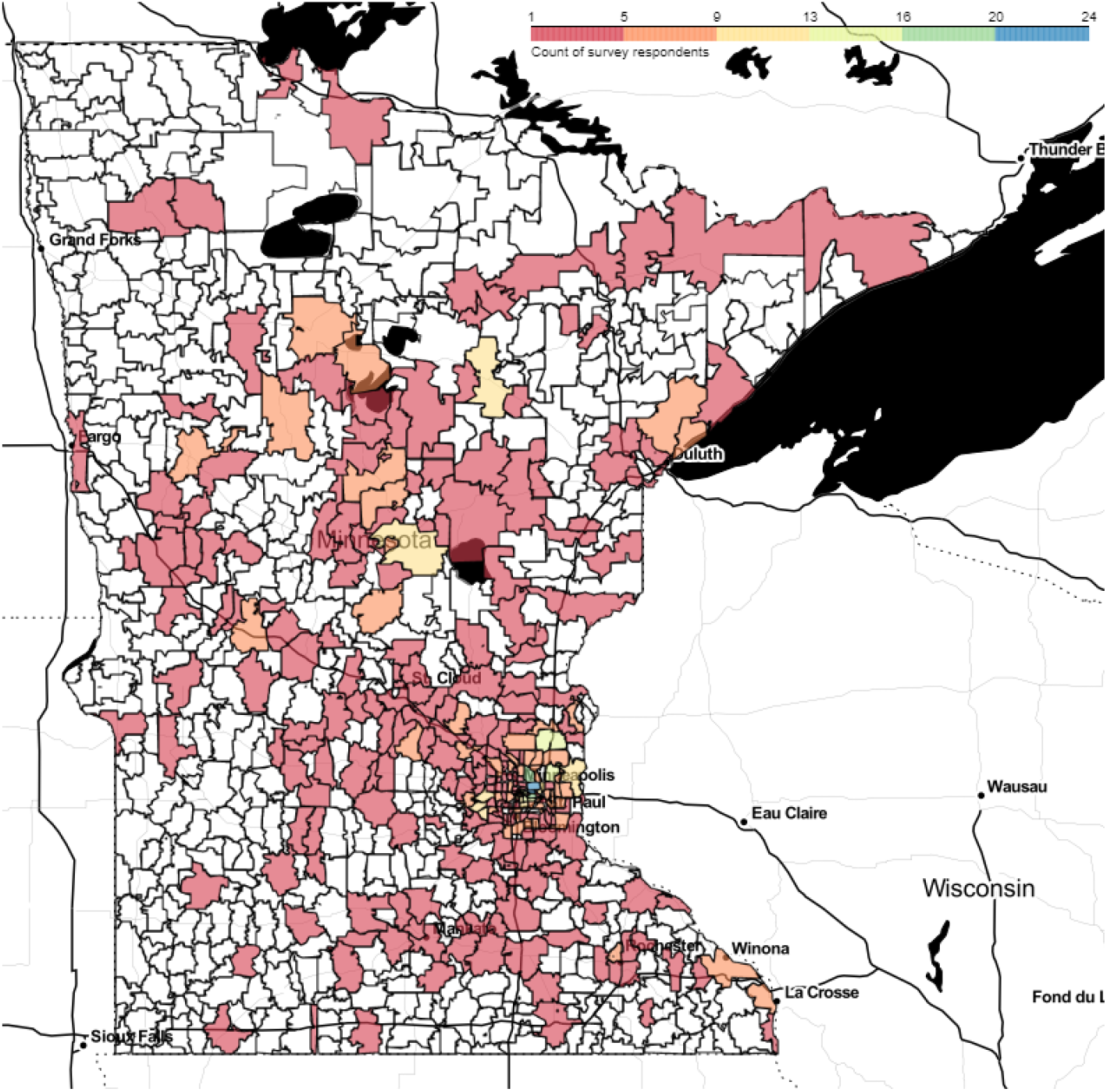
Heatmap of survey responses by zipcode in Minnesota. Color indicates count of survey respondents residing in each zipcode region.

**Supplemental Figure S2.**
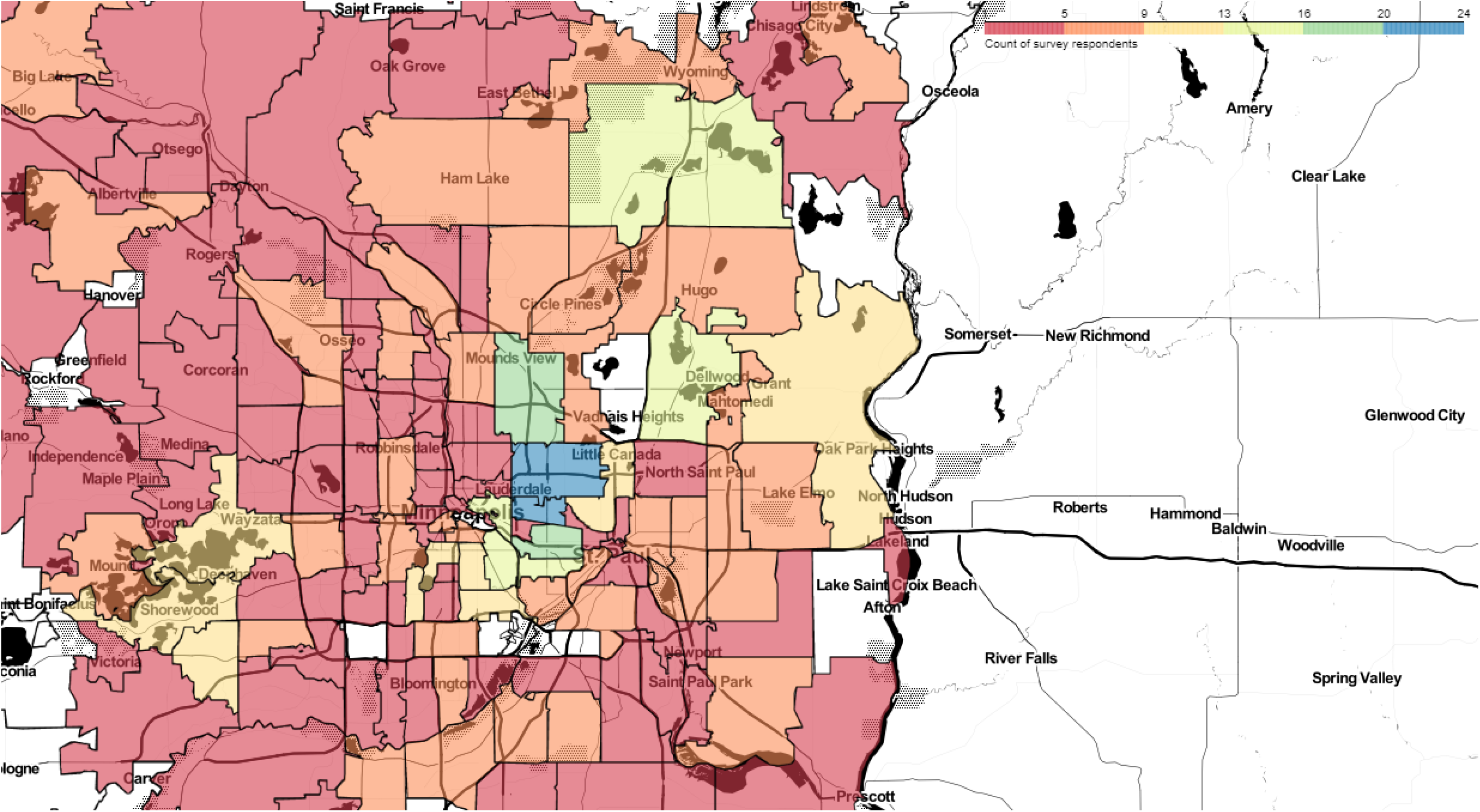
Heatmap of survey responses by zipcode in Minneapolis/Saint Paul area. Color indicates count of survey respondents residing in each zipcode region. The most-represented zipcodes, colored blue in this heatmap, were 55113 (above) and 55108 (below), with 24 respondents residing in each.

## Bibliography

1. Philipp Pietsch, Constanze; Hirsch, editor. Biology and Ecology of Carp. CRC Press, 2015.

2. John D. Koehn. Carp (Cyprinus carpio) as a powerful invader in Australian waterways. doi: 10.1111/j.1365-2427.2004.01232.x.

3. Shin-ichiro S. Matsuzaki, Nisikawa Usio, Noriko Takamura, and Izumi Washitani. Effects of common carp on nutrient dynamics and littoral community composition: roles of excretion and bioturbation. Fundamental and Applied Limnology, 168(1):27–38, 01 2007. doi: 10.1127/1863-9135/2007/0168-0027.

4. Przemyslaw G. Bajer and Peter W. Sorensen. Recruitment and abundance of an invasive fish, the common carp, is driven by its propensity to invade and reproduce in basins that experience winter-time hypoxia in interconnected lakes. Biological Invasions, 12(5):1101–1112, 2010. ISSN 1387-3547. doi: 10.1007/s10530-009-9528-y. Funding Information: Acknowledgments We thank Daryl Ellison, Paul Diedrich, and Lee Sundmark [Minnesota Department of Natural Resources (MN DNR)] for providing historical records of winter hypoxia and fish winterkills in the study lakes. Michael McInerny (MN DNR) and Brett Miller [University of Minnesota (UofMN)] verified age estimates. Mario Travaline (UofMN) conducted carp female egg-counts. Jack Wingate, Tim Cross, Michael McInerny, Nicole Hensel-Welch, Eric Altena (MN DNR), Haude Levesque (UofMN), Tiffany Babich, Greg Aamodt (Carver County Water Management), Dave and Ann Florenzano, John Bushey, and Mike Domke (Lake Riley Association) and other volunteers helped with data collection. Matt Hennen helped with carp telemetry. Ken Seemann conducted seining operations. Paul Brown (Department of Primary Industries, Victoria, Australia) helped with developing procedures to age common carp in our laboratory. Howard Peterson provided encouragement. This research was supported by grants from the Minnesota Environmental and Natural Resources Trust Fund, Riley Purgatory Bluff Creek Watershed District, MN DNR—Ecological Services, Invasive Animals Cooperative Research Centre, and a generous contribution from Bill Oemichen whose early encouragement was greatly appreciated. The Minnesota DNR Fisheries Division also provided in-kind support. We thank an anonymous reviewer for helpful comments on the manuscript.

5. John L. Teem, Luke Alphey, Sarah Descamps, Matt P. Edgington, Owain Edwards, Neil Gemmell, Tim Harvey-Samuel, Rachel L. Melnick, Kevin P. Oh, Antoinette J. Piaggio, J. Royden Saah, Dan Schill, Paul Thomas, Trevor Smith, and Andrew Roberts. Genetic Biocontrol for Invasive Species. Frontiers in Bioengineering and Biotechnology, 8(May):1–18, 2020. ISSN 22964185. doi: 10.3389/fbioe.2020.00452.

6. Peter Martin Grewe. Review and evaluation of the potential of molecular approaches for the environmentally benign management of the common carp (Cyprinus carpio) in Australian waters. Hobart, Tas., CSIRO Div. of Fisheries. Centre for Research on Introduced …, 1996. ISBN 0643059571.

7. Ronald E. Thresher, Keith Hayes, Nicholas J. Bax, John Teem, Tillmann J. Benfey, and Fred Gould. Genetic control of invasive fish: Technological options and its role in integrated pest management. Biological Invasions, 16(6):1201–1216, 2014. ISSN 13873547. doi: 10.1007/s10530-013-0477-0.

8. Bruce L. Webber, S. Raghu, and Owain R. Edwards. Opinion: Is CRISPR-based gene drive a biocontrol silver bullet or global conservation threat? Proceedings of the National Academy of Sciences, 112(34):10565–10567, 2015. ISSN 0027-8424. doi: 10.1073/pnas.1514258112.

9. Paul Slovic. Perception of Risk Author (s): Paul Slovic Published by : American Association for the Advancement of Science Stable URL : http://www.jstor.org/stable/1698637. Advancement Of Science, 236(4799):280–285, 1987.

10. Ronald E Thresher and Armand M Kuris. Options for managing invasive marine species. Biological Invasions, 6(3):295–300, 2004. ISSN 1573-1464.

11. Leah M. Sharpe. Public perspectives on genetic biocontrol technologies for controlling invasive fish. Biological Invasions, 16(6):1241–1256, 2014. ISSN 13873547. doi: 10.1007/s10530-013-0545-5.

12. Cary Funk and Meg Hefferon. Most Americans Accept Genetic Engineering of Animals that Benefits Human Health, but Many Oppose Other Uses. Pew Research Center, 1(July), 2018.

13. Ronald E. Thresher, Michael Jones, and D. Andrew R. Drake. Stakeholder attitudes towards the use of recombinant technology to manage the impact of an invasive species: Sea Lamprey in the North American Great Lakes. Biological Invasions, 21(2):575–586, 2019. ISSN 15731464. doi: 10.1007/s10530-018-1848-3.

14. P. A. Kohl, D. Brossard, D. A. Scheufele, and M. A. Xenos. Public views about editing genes in wildlife for conservation. Conservation Biology, 33(6):1286–1295, 2019. ISSN 15231739. doi: 10.1111/cobi.13310.

15. Michael S. Jones, Jason A. Delborne, Johanna Elsensohn, Paul D. Mitchell, and Zachary S. Brown. Does the U.S. Public support using gene drives in agriculture? And what do they want to know? Science Advances, 5(9), 2019. ISSN 23752548. doi: 10.1126/sciadv.aau8462.

16. Cynthia E. Schairer, Riley Taitingfong, Omar S. Akbari, and Cinnamon S. Bloss. A typology of community and stakeholder engagement based on documented examples in the field of novel vector control. PLoS Neglected Tropical Diseases, 13(11):1–21, 2019. ISSN 19352735. doi: 10.1371/journal.pntd.0007863.

17. Richard A Krueger and M A Casey. A practical guide for applied research. A practical guide for applied research, 2000.

18. United states census bureau quickfacts. https://www.census.gov/quickfacts/fact/table/US/PST045222. Accessed: 2023-03.

19. Airong Zhang, Lucy Carter, Matt Curnock, and Aditi Mankad. Biocontrol of European Carp: Ecological and social risk assessment for the release of Cyprinid herpesvirus 3 (CyHV-3) for carp biocontrol in Australia. CSIRO Land and Water, 3(December), 2019.

20. Everett M Rogers. Attributes of Innovations and Their Rate of Adoption. In Diffusion of Innovations, pages 204–251. The Free Press, NY, 1995.

21. Edith A. MacDonald, Jovana Balanovic, Eric D. Edwards, Wokje Abrahamse, Bob Frame, Alison Greenaway, Robyn Kannemeyer, Nick Kirk, Fabien Medvecky, Taciano L. Milfont, James C. Russell, and Daniel M. Tompkins. Public Opinion Towards Gene Drive as a Pest Control Approach for Biodiversity Conservation and the Association of Underlying Worldviews. Environmental Communication, 14(7):904–918, 2020. ISSN 17524040. doi: 10.1080/17524032.2019.1702568.

22. Mark Damian Duda and Joanne L. Nobile. The fallacy of online surveys: No data are better than bad data. Human Dimensions of Wildlife, 15(1):55–64, 2010. ISSN 10871209. doi: 10.1080/10871200903244250.

23. Mark E. Eiswerth, Steven T. Yen, and G. Cornelis van Kooten. Factors determining awareness and knowledge of aquatic invasive species. Ecological Economics, 70(9):1672–1679, 2011. ISSN 09218009. doi: 10.1016/j.ecolecon.2011.04.012.

24. Lara Hakam. Invasive Species : Public Awareness and Education. Thesis, 2013.

25. Sophie Lecheler and Claes H. de Vreese. How Long Do News Framing Effects Last? A Systematic review of Longitudinal Studies. Annals of the International Communication Association, 40(1):3–30, 2016. ISSN 2380-8985. doi: 10.1080/23808985.2015.11735254.

